# Estimating the Mosquito Density in Guangzhou City, China

**DOI:** 10.1101/2021.11.15.468603

**Authors:** Meili Li, Xian Zhang, Junling Ma

**Affiliations:** College of Science, Donghua University, Shanghai 201620, China; Department of Mathematics and Statistics, University of Victoria, Victoria, BC V8W 2Y2, Canada

**Keywords:** Weather, Mosquito density, Parameters estimation, Markov Chain Monte-Carlo

## Abstract

Mosquito is a vector of many diseases. Predicting the trend of mosquito density is important for early warning and control of mosquito diseases. In this paper, we fit a discrete time mosquito model developed by Gong et al. in 2011, which considers the immature and adult stages, and weather dependent model parameters, to the Breteau Index and Bite Index data for Aedes aegypti in Guangzhou city, China in 2014, as well as the weather data for average temperature, precipitation, evaporation and daylight for the same period. We estimated the model parameters using the Markov Chain Monte-Carlo (MCMC) method. We find that many parameters are not identifiable. We revise and simplify the model so that the parameters of our new model are identifiable. Our results indicate that the model predicted mosquito prevalence agrees well with data. We then use the fitted parameter values against the Breteau Index and Bite Index data for Guangzhou city in 2017 and 2018, and show that the estimated parameter values are applicable for other seasons.

## 1 Introduction

Mosquito is a vector for many diseases, such as dengue fever, malaria, and filariasis etc. Dengue fever, in particular, causes a severe burden globally. It exists in 128 countries (Brady et al., 2012), puts 4 billion people at risk (Bhatt et al., 2013), and is estimated to infect 390 million people per year (Waggoner et al., 2016). Aedes aegypti is the primary vector of Dengue fever. Estimating the trend of mosquito density is crucial to give early warnings to an outbreak.

Many countries and regions publish mosquito surveillance data, e.g., the Breteau Index (BI) and Bite Index. However, to better predict mosquito prevalence, we need to understand how these indices are related to the mosquito population dynamics. Mosquito population dynamics is heavy influenced by weather conditions (Brower, 2001; Githeko et al., 2000; Nagao et al., 2003; Tanser et al., 2003). Several models have been developed to estimate mosquito density by incorporating weather factors (Chaves et al., 2012; Gong et al., 2011; Jia et al., 2016; Otero et al., 2006; Tran et al., 2013; Wang et al., 2016; Wong et al., 2011). Jia et al. (2016) proposed a new climate-driven model of Aedes albopictus suitable for most Ae. albopictus-colonized areas. The new model was an improvement of the previous work by quantifying the conditions when the diapause arose and adjusting the model parameters, such as development rates, mortality rates and fecundity rates. Then the new model was validated with the entomological field data for Guangzhou city and Shanghai, China. The results indicated that the improvement was significant over the basic model. Gong et al. (2011) established a climate-based model for West Nile Culex Mosquito Vectors in the Northeastern US in 2011, based on the development results from laboratory studies. The model was optimized by a parameter-space search within biological bounds. The simulated abundance was highly correlated with actual mosquito numbers and validated with field data. Wang et al. (2016) modified the Gong et al. (Gong et al., 2011) model, and fitted the model to the 2014 and 2015 BI data using the least squares method, estimated key model parameters, and validated their estimation using the BI data for 3 districts in Guangzhou city.

However, some key questions have not been addressed by these studies. What are the uncertainties of the estimated parameters? Are the estimated parameters applicable for other seasons?

In this paper, we start by fitting the discrete time Gong et al. (Gong et al., 2011) model to the BI and adult mosquito density (Bite Index) data for 12 districts for Guangzhou city in 2014, as well as the weather data including temperature, precipitation, evaporation and daylight for Guangzhou city in the same period. We will determine the unidentifiable parameters, and revise the model accordingly so that the model parameters are identifiable. We will use the Markov Chain Monte-Carlo method for model fitting, to estimate the posterior distributions of the parameters. At last, we will apply the model parameters to the BI and Bite Index data for Guangzhou city in 2017 and 2018, to determine whether the fitted parameters are application to other seasons.

## 2 Methods

### 2.1 Model

We use the Gong et al. (Gong et al., 2011) model for the mosquito population dynamics, which is a discrete time model with the immature (*J*, including egg, larva and pupa) and adult (*A*) stages of mosquitoes,

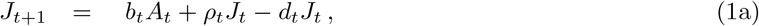

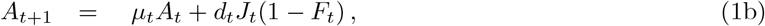

where the fecundity rate *b*_*t*_ is assumed to exponentially increase with the moisture index *M*_*t*_, i.e.,

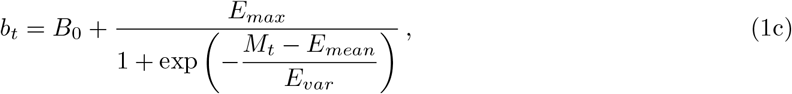

and the moisture index *M*_*t*_ is a 7-day sum of the difference between precipitation *P*_*t*_ (in mm) and evaporation *E*_*t*_ (in mm) for seven consecutive days (which is proportional to the moving average), namely,

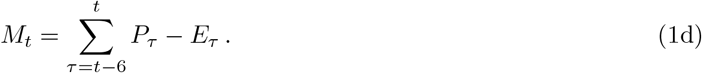

The meanings of the parameters *E*_*max*_, *E*_*mean*_, *E*_*var*_ (and others will be introduced later) are listed in Table 1 The survival rate of the immature mosquitoes *ρ*_*t*_ is assumed to depend on the temperature *T*_*t*_ (in degree Celsius) as a bell curvev, i.e.,

**Table 1:** Definitions of the parameters used in Model (1) Parameters Definition (Units)

| Parameters | Definition (Units) |
| --- | --- |
| $B_0$ | Minimum baseline fecundity rate. |
| $E_{max}$ | Maximum fecundity rate above baseline. |
| $E_{mean}$ | Value at which the moisture index produces 50% of $E_{max}$ (mm). |
| $E_{var}$ | Variance of function. |
| $B_1$ | Maturation rate assuming no temperature inactivation of the critical enzyme. |
| $H_A$ | Enthalpy of activation of the reaction that is catalyzed by the enzyme ($cal\ mol^{-1}$). |
| $H_H$ | Change of Enthalpy with high temperature inactivation of the enzyme ($cal\ mol^{-1}$). |
| $T_H$ | Temperature where 50% of the enzyme is inactivated by high temperature ($^{\circ}C$). |
| $\mu_1$ | Survival rate at optimal temperature for immatures. |
| $T_{01}$ | Optimal temperature for survival of immatures ($^{\circ}C$). |
| $v_1$ | Variance of function. |
| $\mu_2$ | Survival rate at optimal temperature for adults. |
| $T_{02}$ | Optimal temperature for survival of adults ($^{\circ}C$). |
| $v_2$ | Variance of function. |
| $k$ | Decay rate of diapausing rate with day length. |

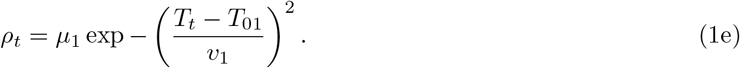

The maturation rate of immature mosquitoes *d*_*t*_ is assumed to be a slightly skewed single humped curve of the temperature *T*_*t*_, i.e.,

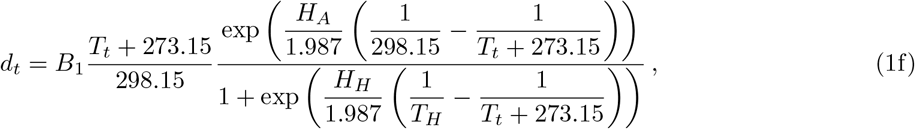

based on the Sharpe and DeMichele equation (Rueda et al., 1990), where the constants are determined empirically. The survival rate of the adults *μ*_*t*_ is assumed to be a bell curve of the temperature *T*_*t*_, i.e.,

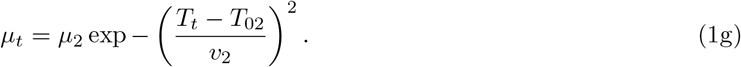

The diapause rate of the adults *F*_*t*_ is assumed to be a linear function of the day length *H*_*t*_ (in hours), i.e.,

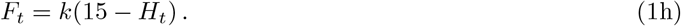

### 2.2 Mosquito density data

The mosquito monitoring data for Guangzhou city were obtained from the Guangzhou Center for Disease Control and Prevention (GZCDC) (Guangzhou Center for Disease Control and Prevention, 2021). The data includes the daily Breteau Index and Bite Index for Aedes aegypti for the 12 districts in Guangzhou city, China for the period of September 26 to November 3, 2014, and weekly Breteau Index and Bite Index for the periods August 28 – December 17, 2017 and July 30 – October 15, 2018. These indices are not the real densities, but are density indicators of immatures and adults respectively. We use the 2014 data for model fitting, and the 2017 and 2018 data for model validation.

Note that, before February 2014, there were 12 districts in Guangzhou city. In February 2014, the Luogang district was merged into the Huangpu district in Guangzhou, reducing the number of districts to 11. But the 2014 mosquito data were still published for the original 12 districts. Since 2015, the mosquito data have been published for the 11 new districts.

### 2.3 Weather data

The temperature, precipitation and evaporation data for Guangzhou city were obtained from the network (Chinese Software Develop Net, 2020), and the daylight data for Guangzhou city were downloaded from the network (Convenient Inquiry Network, 2021).

### 2.4 Parameter estimation

To link the BI and Bite Index to immature and adult mosquito densities, we let *p*_*J*_ to be the capture probability of immature mosquitoes, and *p*_*A*_ to be the capture probability of adults. Therefore, expected BI is equal to *J*_*t*_*p*_*J*_ and the expected Bite Index is *A*_*t*_*p*_*A*_. Since the model is a linear system, a constant multiple of the solution is still a solution. Thus we cannot individually estimate both *p*_*A*_ and *p*_*J*_. Instead, we estimate their ratio

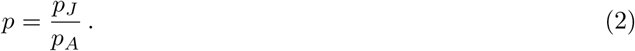

The population density is related to counts, which typically assumed to be a Poisson random variable and has a variance similar to the mean. However, the BI *X*_*t*_ and the Bite Index *Y*_*t*_ are not true counts. So, we assume that they are normally distributed with a mean *A*_*t*_ and variance *A*_*t*_ for the adults, and a mean *J*_*t*_*p* and vairance *J*_*t*_*p* for the immatures, i.e.,

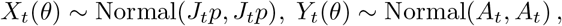

where the parameter

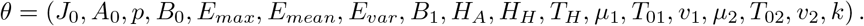

We fit the model to the mosquito and weather data for the 12 districts in Guangzhou city in 2014 individually. For each district, the initial densities *J*_0_ and *A*_0_, and the ratio *p* of the capture probabilities, are estimated individually, while the other parameters are assumed to be the same across districts. We use the Markov Chain Monte-Carlo (MCMC) method with Gibbs sampling via the *R2jags* package in R.

We used wide uninformative uniform prior distributions for the fitting. The prior distributions are listed in Table 2. We used 4 chains, 10,000 iterations with the first 6,000 iterations discarded as a burn-in period.

**Table 2:** The prior distributions of parameters for the 12 districts in Guangzhou city based on the basic model.

| Parameters | Prior | Parameters | Prior | Parameters | Prior |
| --- | --- | --- | --- | --- | --- |
| $J_0$ (Yuexiu District) | $U(0, 120)$ | $B_0$ | $U(0, 5)$ | $p$ (Yuexiu District) | $U(0, 1)$ |
| $A_0$ (Yuexiu District) | $U(0, 20)$ | $E_{max}$ | $U(0, 60)$ | $p$ (Haizhu District) | $U(0, 1)$ |
| $J_0$ (Haizhu District) | $U(0, 300)$ | $E_{mean}$ | $N(0, 30)$ | $p$ (Huangpu District) | $U(0, 1)$ |
| $A_0$ (Haizhu District) | $U(0, 40)$ | $E_{var}$ | $U(0, 40)$ | $p$ (Tianhe District) | $U(0, 1)$ |
| $J_0$ (Huangpu District) | $U(0, 1800)$ | $B_1$ | $U(0, 1)$ | $p$ (Baiyun District) | $U(0, 1)$ |
| $A_0$ (Huangpu District) | $U(0, 30)$ | $H_A$ | $U(0, 70000)$ | $p$ (Liwan District) | $U(0, 1)$ |
| $J_0$ (Tianhe District) | $U(0, 300)$ | $H_H$ | $U(0, 210000)$ | $p$ (Nansha District) | $U(0, 2)$ |
| $A_0$ (Tianhe District) | $U(0, 30)$ | $T_H$ | $U(0, 400)$ | $p$ (Panyu District) | $U(0, 1)$ |
| $J_0$ (Baiyun District) | $U(0, 400)$ | $\mu_1$ | $U(0, 1)$ | $p$ (Luogang District) | $U(0, 1)$ |
| $A_0$ (Baiyun District) | $U(0, 50)$ | $T_{01}$ | $U(0, 40)$ | $p$ (Huadu District) | $U(0, 1)$ |
| $J_0$ (Liwan District) | $U(0, 1600)$ | $v_1$ | $U(5, 160)$ | $p$ ((Zengcheng City) | $U(0, 1)$ |
| $A_0$ (Liwan District) | $U(0, 80)$ | $\mu_2$ | $U(0, 1)$ | $p$ (Conghua City) | $U(0, 1)$ |
| $J_0$ (Nansha District) | $U(0, 350)$ | $T_{02}$ | $U(10.80)$ | | |
| $A_0$ (Nansha District) | $U(0, 30)$ | $v_2$ | $U(5, 200)$ | | |
| $J_0$ (Panyu District) | $U(0, 700)$ | $k$ | $U(0, 0.2)$ | | |
| $A_0$ (Panyu District) | $U(0, 40)$ | | | | |
| $J_0$ (Luogang District) | $U(0, 1000)$ | | | | |
| $A_0$ (Luogang District) | $U(0, 50)$ | | | | |
| $J_0$ (Huadu District) | $U(0, 300)$ | | | | |
| $A_0$ (Huadu District) | $U(0, 200)$ | | | | |
| $J_0$ (Zengcheng City) | $U(0, 80)$ | | | | |
| $A_0$ (Zengcheng City) | $U(0, 50)$ | | | | |
| $J_0$ (Conghua City) | $U(0, 100)$ | | | | |
| $A_0$ (Conghua City) | $U(0, 100)$ | | | | |

## 3 Results

The posterior distributions for the initial conditions for the 12 districts, model parameters for all districts, and the ratio of capture probabilities for the 12 districts, are shown in Figure 1, Figure 2 and Figure 3 respectively. The parameters *B*_1_, *H*_*H*_, *H*_*A*_, *T*_*H*_ related to the maturation rate *d*_*t*_, and the *k* of the diapause rate *F*_*t*_ cannot be identified, as their posterior distributions are uniform even when increasing the widths of the prior distribution. In addition, the parameter *v*_2_ of the adult survival rate *μ*_*t*_ is also unidentifiable because its posterior density kept rising even when increasing the widths of the prior distribution.

**Figure 1:**
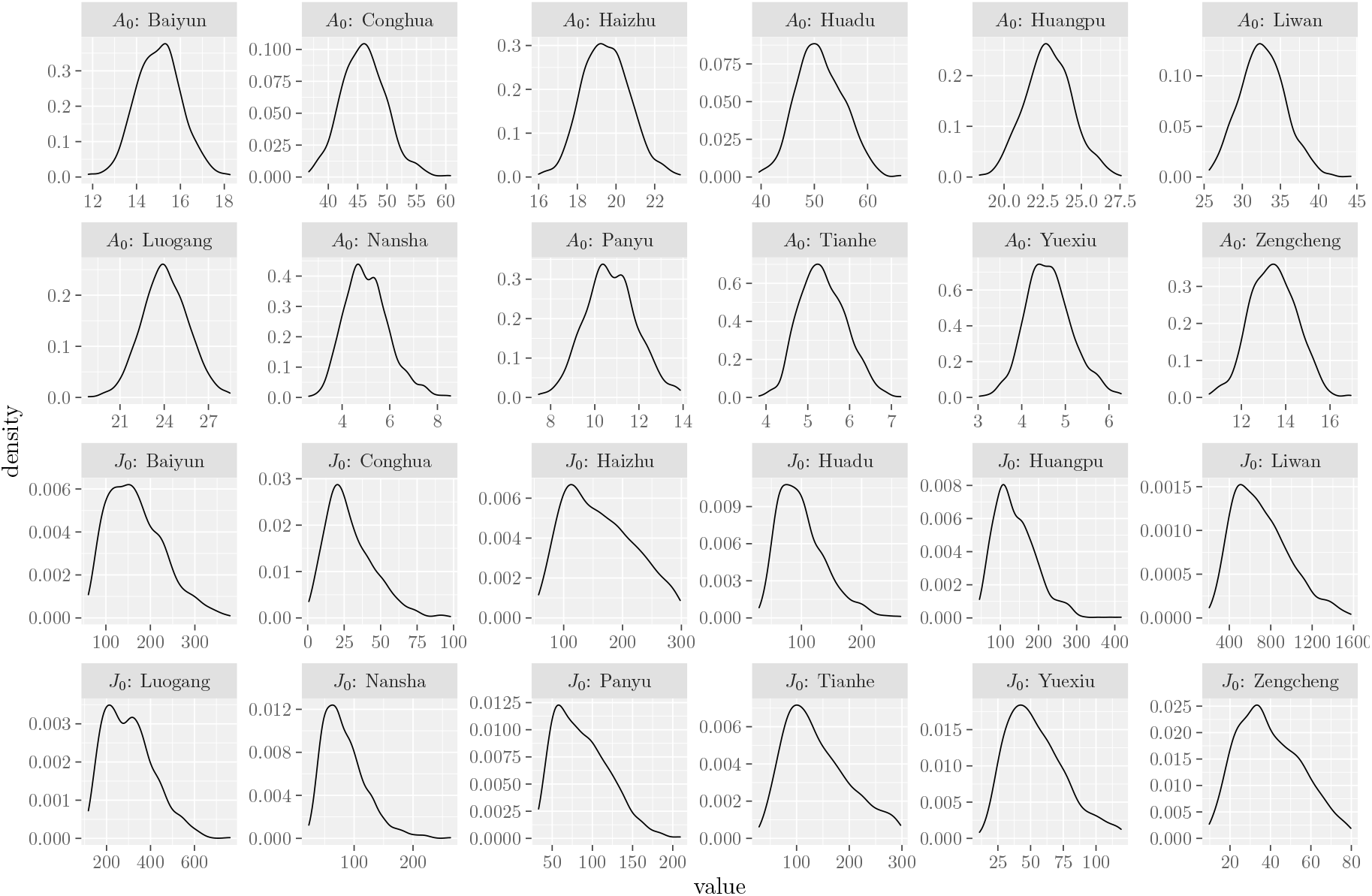
The posterior distributions of initial values of Model (1) for the 12 districts in Guangzhou city.

**Figure 2:**
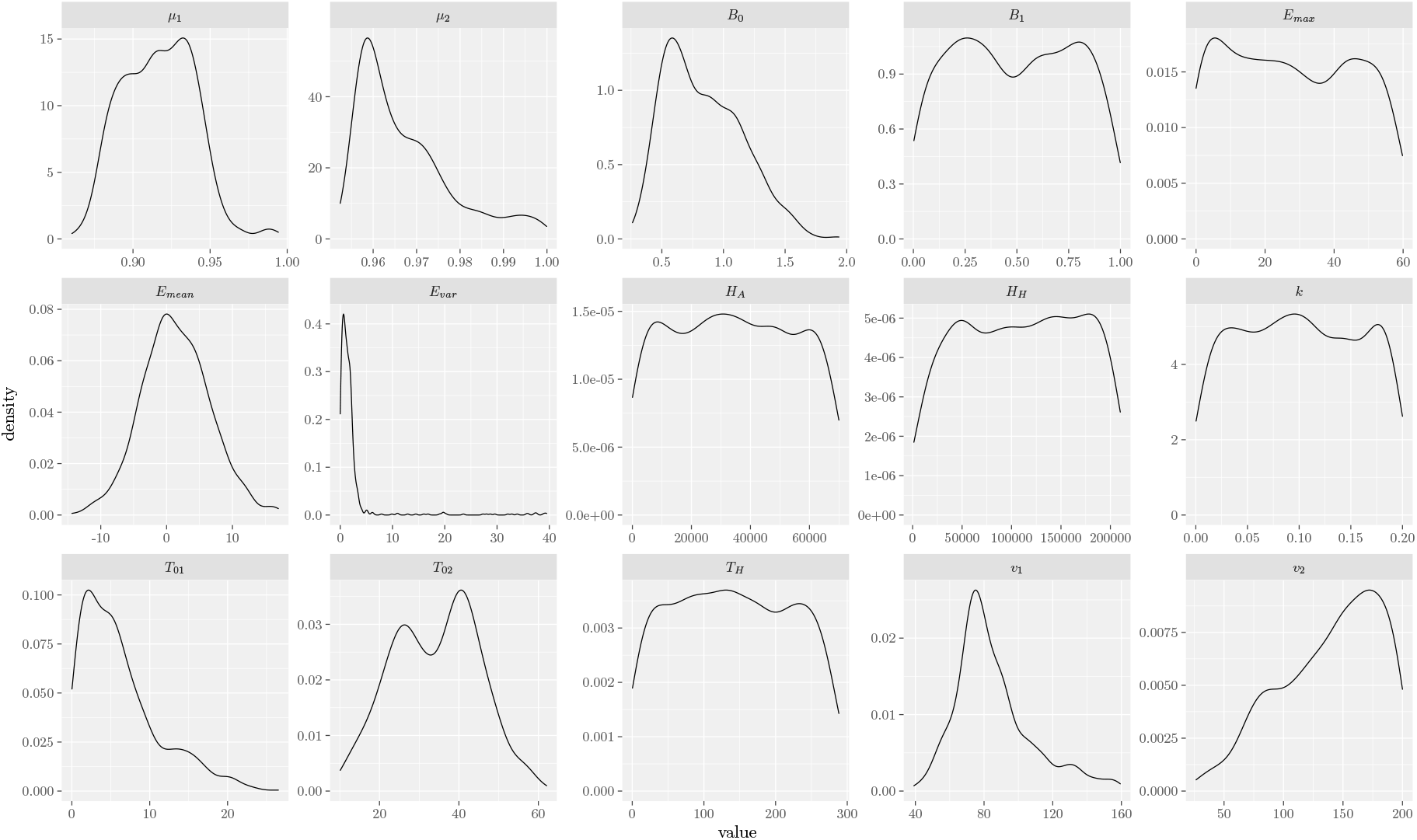
The posterior distributions of the parameters of Model (1).

**Figure 3:**
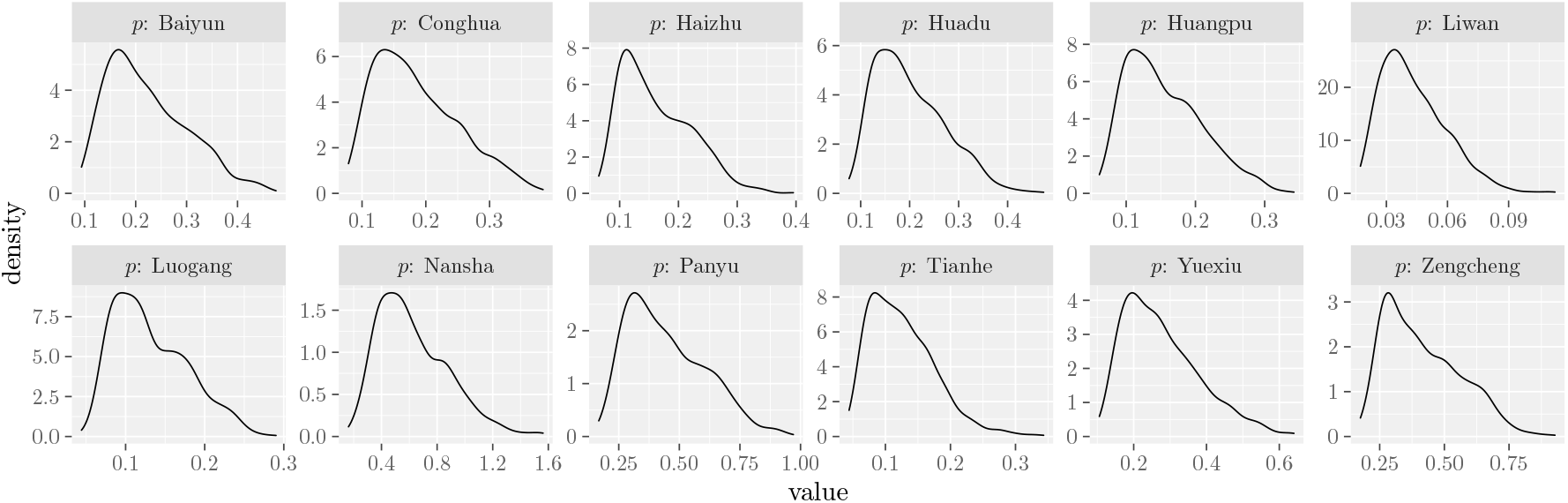
The posterior distributions of capture probability ratios of Model (1) for the 12 districts in Guangzhou city.

### A new maturation rate

The maturation rate *d*_*t*_ defined in (1f) is a single humped function of the temperature *T*_*t*_. For simplicity, we replace it with a bell curve:

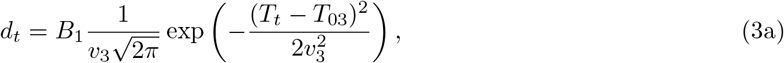

where *T*_03_ is the peak and *v*_3_ is the spread of the bell curve.

### A new adult survival rate

From (1g), the adult survival rate is approximately a constant (independent of the temperature *T*_*t*_) if *v*_2_ becomes very large, which is suggested by our fitting results. Thus, we assume that

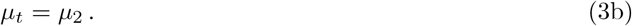

### A new diapause rate

Based on the work reported by Spielman (Spielman, 2001), the diapause rate was a function of decreasing hours of daylight *H*_*t*_, ranging from 0 to 1. We assume that it decreases exponentially with the daylight *H*_*t*_,

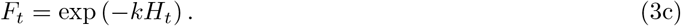

### The simulation results of improved model

We fitted the basic model (1)(a,b) with (3)(a,b,c) to the same data. All the parameters are identifiable. The prior distributions of parameters were shown in Table 3. The posterior distributions are given in Figure 4, Figure 5 and Figure 6, the mean and the confidence intervals of the model parameters (excluding the initial densities and the ratio of the capture probabilities) are listed in Table 4.

**Table 3:** The prior distributions of parameters for the 12 districts in Guangzhou city based on the improved model.

| Parameters | Prior | Parameters | Prior | Parameters | Prior |
| --- | --- | --- | --- | --- | --- |
| $J_0$ (Yuexiu District) | $U(0.50)$ | $B_0$ | $U(0, 5)$ | $p$ (Yuexiu District) | $U(0, 2)$ |
| $A_0$ (Yuexiu District) | $U(0, 20)$ | $E_{max}$ | $U(0, 30)$ | $p$ (Haizhu District) | $U(0, 2)$ |
| $J_0$ (Haizhu District) | $U(0, 100)$ | $E_{mean}$ | $U(0, 30)$ | $p$ (Huangpu District) | $U(0, 1)$ |
| $A_0$ (Haizhu District) | $U(0, 40)$ | $E_{var}$ | $U(0, 30)$ | $p$ (Tianhe District) | $U(0, 1)$ |
| $J_0$ (Huangpu District) | $U(0, 2000)$ | $B_1$ | $U(0, 1)$ | $p$ (Baiyun District) | $U(0, 1)$ |
| $A_0$ (Huangpu District) | $U(0, 30)$ | $T_{03}$ | $U(0, 40)$ | $p$ (Liwan District) | $U(0, 1)$ |
| $J_0$ (Tianhe District) | $U(0, 250)$ | $v_3$ | $U(0, 10)$ | $p$ (Nansha District) | $U(0, 1)$ |
| $A_0$ (Tianhe District) | $U(0, 30)$ | $\mu_1$ | $U(0, 1)$ | $p$ (Panyu District) | $U(0, 0.8)$ |
| $J_0$ (Baiyun District) | $U(0, 400)$ | $\mu_2$ | $U(0, 0.95)$ | $p$ (Luogang District) | $U(0, 1)$ |
| $A_0$ (Baiyun District) | $U(0, 50)$ | $T_{01}$ | $U(0, 40)$ | $p$ (Huadu District) | $U(0, 3.5)$ |
| $J_0$ (Liwan District) | $U(0, 1500)$ | $v_1$ | $U(0, 60)$ | $p$ (Zengcheng City) | $U(0, 5.5)$ |
| $A_0$ (Liwan District) | $U(0, 60)$ | $k$ | $U(0, 0.1)$ | $p$ (Conghua City) | $U(0, 5)$ |
| $J_0$ (Nansha District) | $U(0, 350)$ | | | | |
| $A_0$ (Nansha District) | $U(0, 30)$ | | | | |
| $J_0$ (Panyu District) | $U(0, 700)$ | | | | |
| $A_0$ (Panyu District) | $U(0, 40)$ | | | | |
| $J_0$ (Luogang District) | $U(0, 1000)$ | | | | |
| $A_0$ (Luogang District) | $U(0, 50)$ | | | | |
| $J_0$ (Huadu District) | $U(0, 300)$ | | | | |
| $A_0$ (Huadu District) | $U(0, 200)$ | | | | |
| $J_0$ (Zengcheng City) | $U(0, 60)$ | | | | |
| $A_0$ (Zengcheng City) | $U(0, 50)$ | | | | |
| $J_0$ (Conghua City) | $U(0, 100)$ | | | | |
| $A_0$ (Conghua City) | $U(0, 100)$ | | | | |

**Table 4:** The estimated parameters values (with 95% confidence intervals in parenthesizes).

| Parameters | Values | Parameters | Values |
| --- | --- | --- | --- |
| $B_0$ | 0.038(0.017,0.087) | $v_3$ | 0.33(0.22,0.46) |
| $E_{max}$ | 13.40 (0.08,29.10) | $\mu_1$ | 0.997(0.99,1) |
| $E_{mean}$ | 1.72 (-7.98,11.78) | $\mu_2$ | 0.93(0.92,0.93) |
| $E_{var}$ | 1.44 (0.043,3.94) | $T_{01}$ | 20.94(18.91,22.62) |
| $B_1$ | 0.22 (0.18,0.26) | $v_1$ | 31.04(22.21,42.63) |
| $T_{03}$ | 23.02 (22.91,23.21) | $k$ | 0.005(0.002,0.01) |

**Figure 4:**
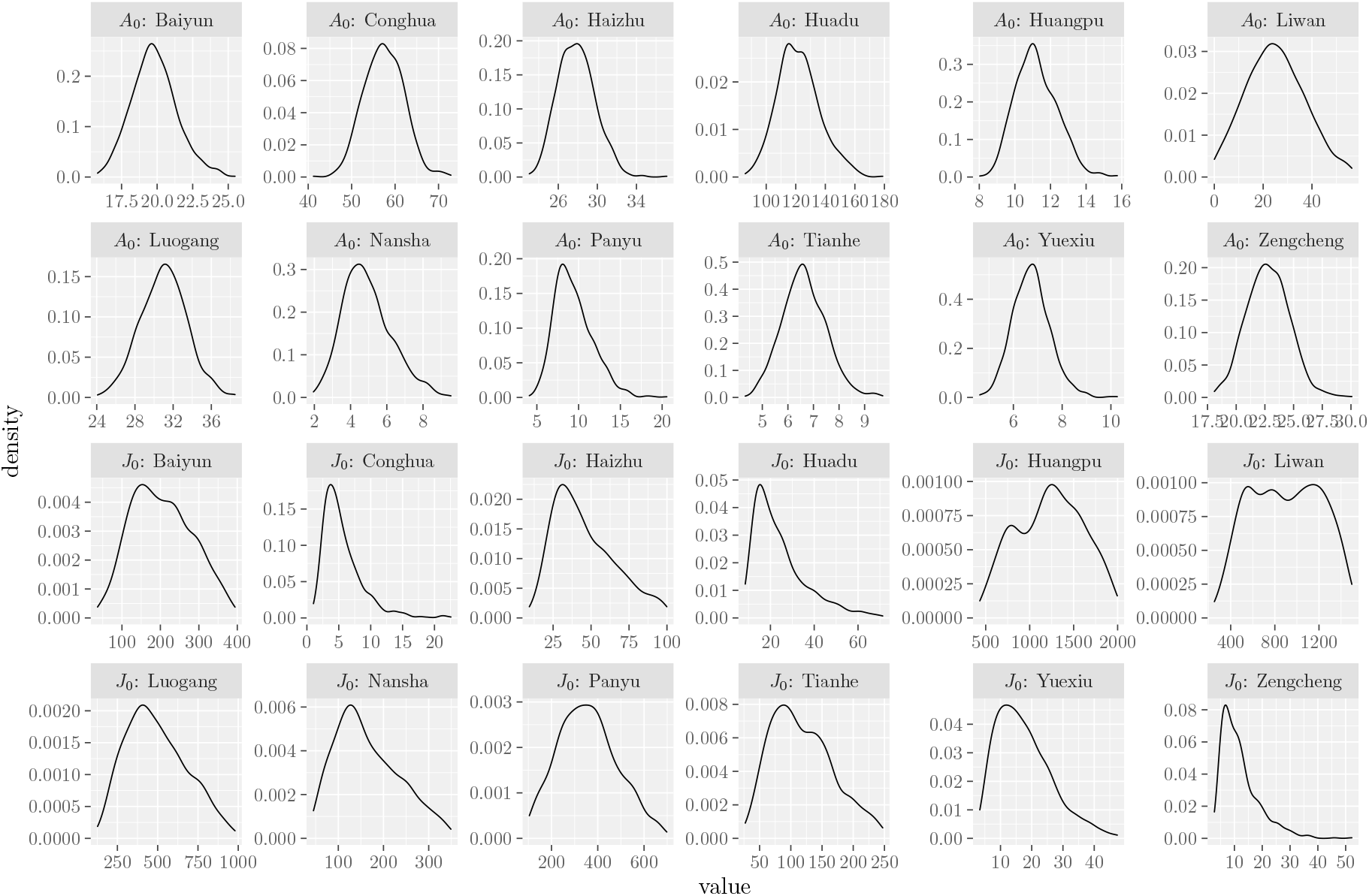
The posterior distributions of initial values of Model (3) for the 12 districts in Guangzhou city.

**Figure 5:**
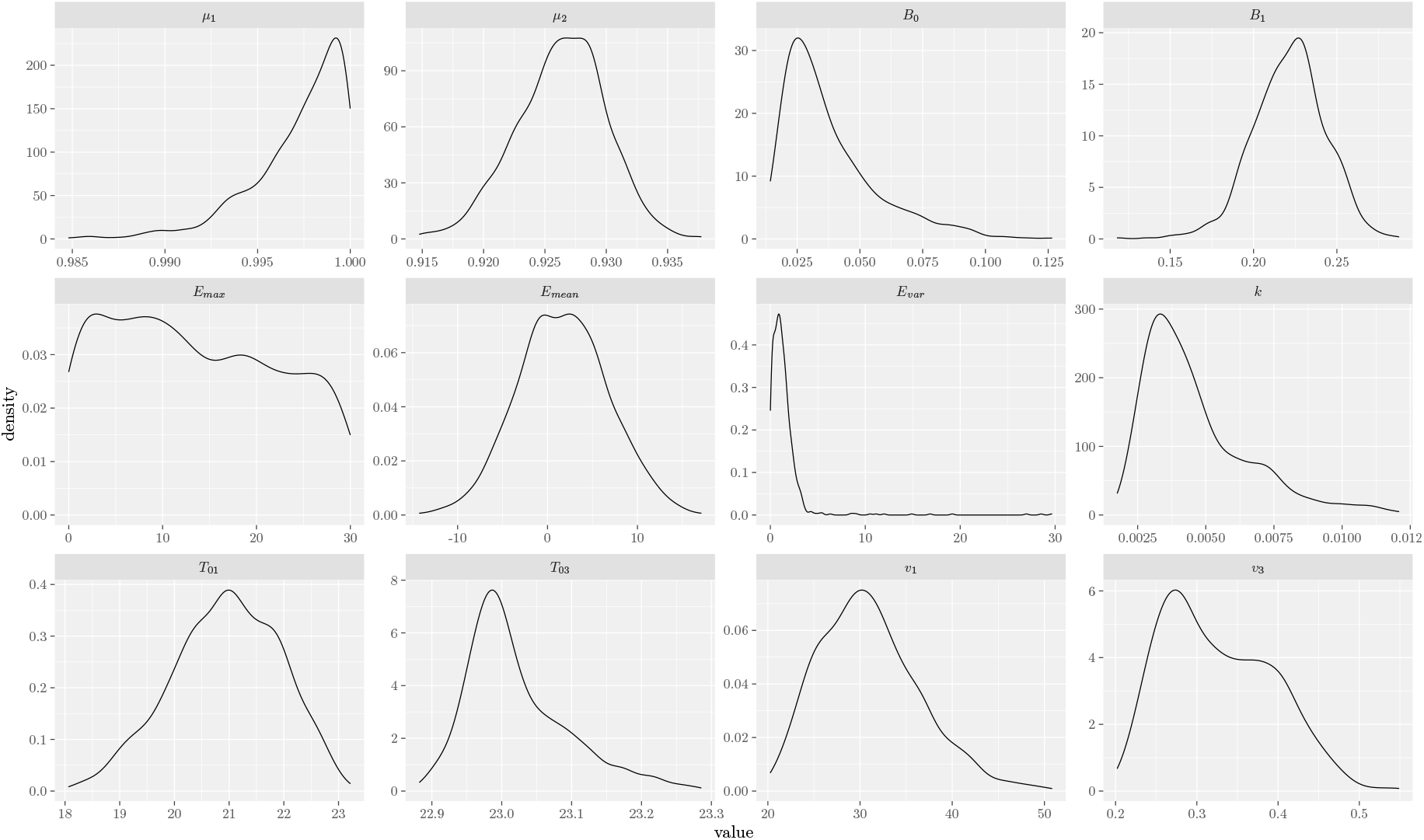
The posterior distributions of parameters of Model (3).

**Figure 6:**
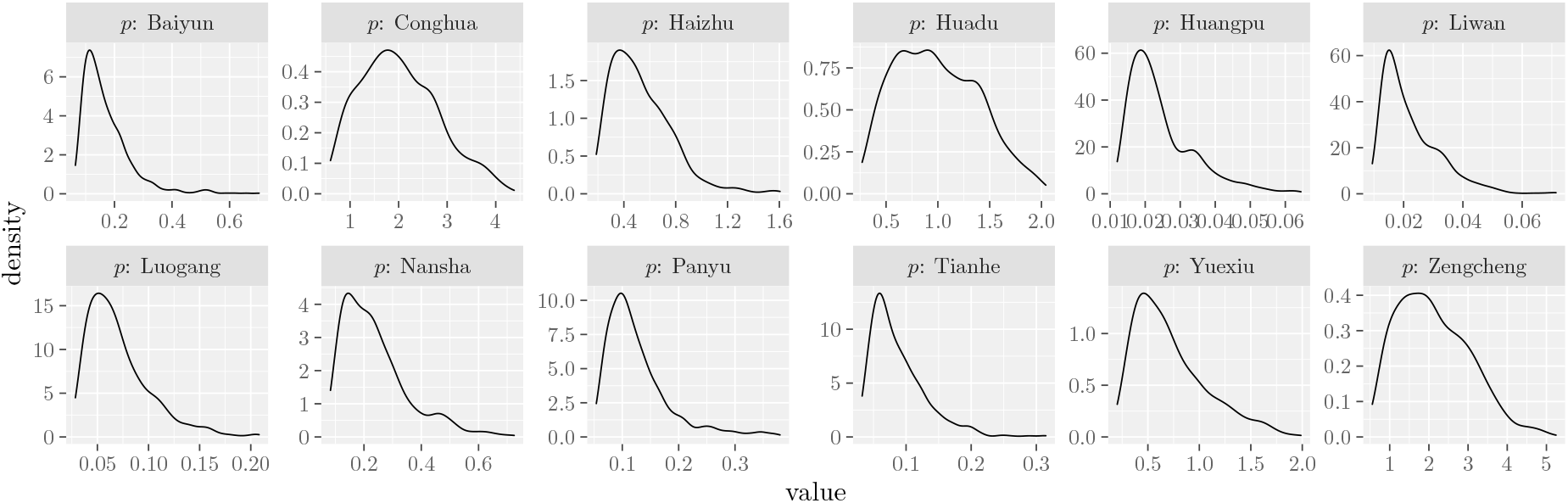
The posterior distributions of capture probability ratios of Model (3) for the 12 districts in Guangzhou city.

In Figure 7, we plot the predicted mosquito density using the model (1)(a,b) with (3)(a,b,c) and the point estimates (the mean of the posterior distributions) of the fitted parameters, and compare with the BI and Bite Index data (green dots) for the 12 districts in Guangzhou city. The correspondence between the simulation result and the data is very good.

**Figure 7:**
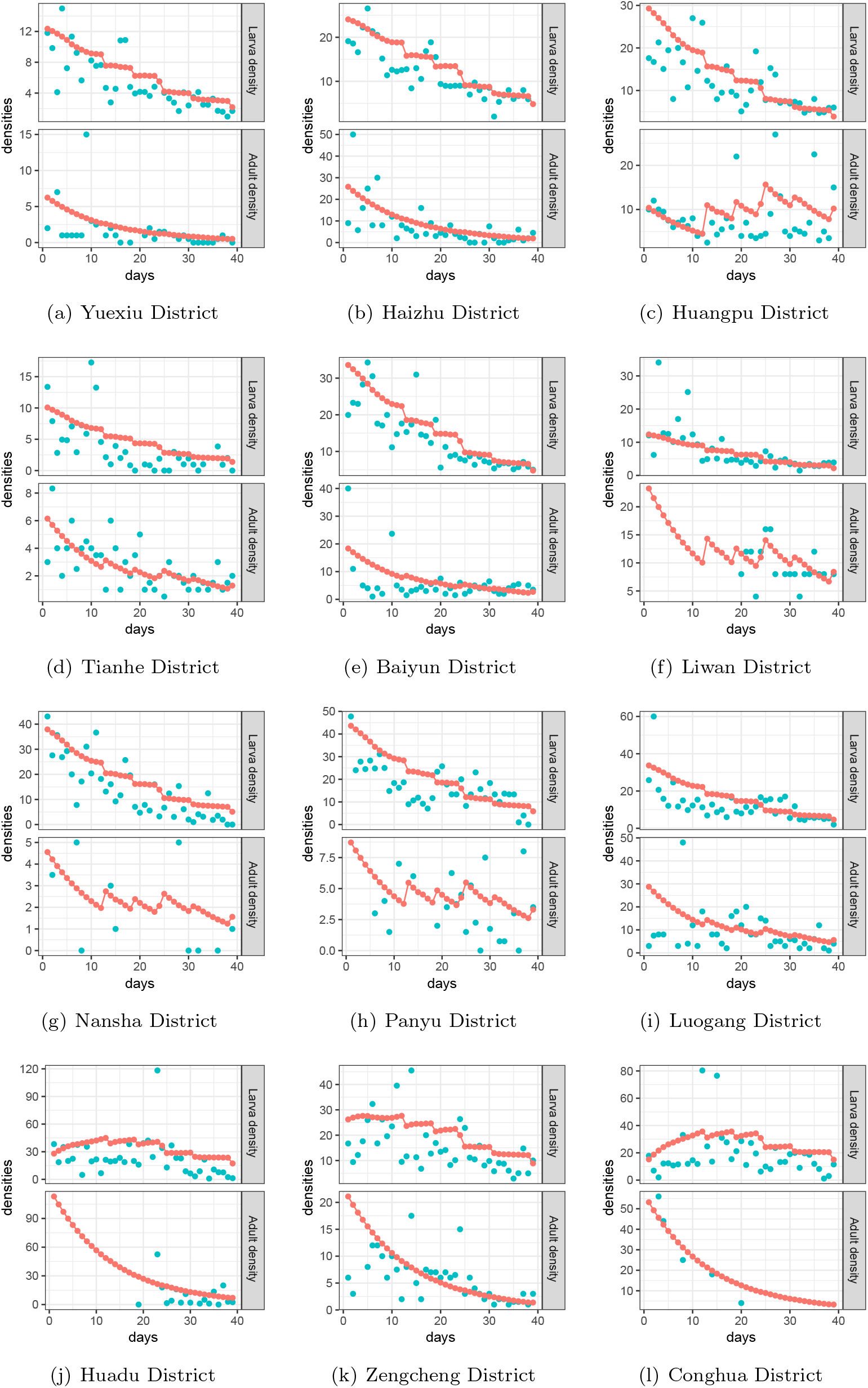
The comparison of the predicted mosquito density (red dots) with the Breteau Index and Bite Index for 12 districts in Guangzhou city in 2014 (blue dots).

## 4 Model validation

Are the model parameters estimated in the previous section applicable to other years? To answer this questions, we fit our model (1)(a,b) with (3)(a,b,c) to the mosquito surveillance data for Guangzhou city in 2017 and 2018. The initial densities *J*_0_ and *A*_0_, and the ratio of the capturing probability *p*, are estimated separately for each district, while the other estimated parameters are listed in Table 4.

The estimated initial values and capture probability ratios for 2017 are shown in Table 5, and the comparisons between the model predictions and BI and Bite Index data for the 11 districts in Guangzhou city are shown in Figure 8. For 2018, the fitted parameter values are summarized in Table 6, and the comparisons of model predictions and data are shown in Figure 9. Both figures show that the simulated abundance better predicts the observed mosquito density data. Thus, the parameter values estimated for the model (1)(a,b) with (3)(a,b,c) for 2014 are also suitable for 2017 and 2018.

**Table 5:** The estimated values of initial values and capture probability ratios for the 11 districts in Guangzhou city in 2017 (with 97.5% confidence intervals in parenthesizes).

| Parameters | Values | Parameters | Values |
| --- | --- | --- | --- |
| $J_0$ (Yuexiu District) | 56.755 (16.122,118.365) | $J_0$ (Nansha District) | 16.341 (5.689,38.635) |
| $A_0$ (Yuexiu District) | 1.502 (0.956,2.225) | $A_0$ (Nansha District) | 0.866(0.527,1.316) |
| $J_0$ (Haizhu District) | 113.04 (42.164,203.553) | $J_0$ (Panyu District) | 31.17 (11.1,74.699) |
| $A_0$ (Haizhu District) | 1.836 (1.059,2.829) | $A_0$ (Panyu District) | 2.902 (2.126,3.824) |
| $J_0$ (Huangpu District) | 3.627(0.895,10.515) | $J_0$ (Huadu District) | 11.683(3.861,29.16) |
| $A_0$ (Huangpu District) | 0.532 (0.268,0.984) | $A_0$ (Huadu District) | 1.179 (0.775,1.722) |
| $J_0$ (Tianhe District) | 101.659 (31.521,194.557) | $J_0$ (Zengcheng District) | 43.288 (10.589,142.601) |
| $A_0$ (Tianhe District) | 1.704 (0.978,2.634) | $A_0$ (Zengcheng District) | 2.817(1.511,3.958) |
| $J_0$ (Baiyun District) | 0.985(0.018,4.056) | $J_0$ (Conghua District) | 7.712(1.3,27.931) |
| $A_0$ (Baiyun District) | 1.081(0.703,1.589) | $A_0$ (Conghua District) | 1.222(0.771,1.802) |
| $J_0$ (Liwan District) | 12.689 (3.802,33.157) | $p$ (Liwan District) | 0.089(0.037,0.177) |
| $A_0$ (Liwan District) | 0.584 (0.309,1.022) | $p$ (Nansha District) | 0.095 (0.043,0.174) |
| $p$ (Yuexiu District) | 0.013 (0.006,0.027) | $p$ (Panyu District) | 0.049(0.028,0.075) |
| $p$ (Haizhu District) | 0.019(0.01,0.036) | $p$ (Huadu District) | 0.131(0.065,0.212) |
| $p$ (Huangpu District) | 0.207(0.088,0.382) | $p$ (Zengcheng District) | 0.045 (0.02,0.073) |
| $p$ (Tianhe District) | 0.009 (0.004,0.018) | $p$ (Conghua District) | 0.077(0.038,0.132) |
| $p$ (Baiyun District) | 0.175 (0.099,0.274) | | |

**Table 6:** The estimated values of initial values and capture probability ratios for the 11 districts in Guangzhou city in 2018 (with 97.5% confidence intervals in parenthesizes).

| Parameters | Values | Parameters | Values |
| --- | --- | --- | --- |
| $J_0$ (Yuexiu District) | 68.93 (3.342,196.729) | $J_0$ (Nansha District) | 7.317(0.155,29.086) |
| $A_0$ (Yuexiu District) | 6.541(5.383,7.923) | $A_0$ (Nansha District) | 9.793 (7.527,10.233) |
| $J_0$ (Haizhu District) | 10.166 (0.332,37.422) | $J_0$ (Panyu District) | 9.266 (0.265,31.315) |
| $A_0$ (Haizhu District) | 5.456(4.405,6.68) | $A_0$ (Panyu District) | 7.196 (5.997,8.547) |
| $J_0$ (Huangpu District) | 15.631 (0.52,59.097) | $J_0$ (Huadu District) | 10.626 (0.328,38.59) |
| $A_0$ (Huangpu District) | 11.611 (10.029,13.479) | $A_0$ (Huadu District) | 4.799 (3.789,5.928) |
| $J_0$ (Tianhe District) | 8.95 (0.303,35.068) | $J_0$ (Zengcheng District) | 11.623(0.321,39.162) |
| $A_0$ (Tianhe District) | 2.577 (1.898,3.432) | $A_0$ (Zengcheng District) | 5.079 (4.041,6.199) |
| $J_0$ (Baiyun District) | 18.262(0.42,65.776) | $J_0$ (Conghua District) | 893.886 (558.038,1373.509) |
| $A_0$ (Baiyun District) | 5.586 (4.511,6.878) | $A_0$ (Conghua District) | 0.368 (0.16,0.759) |
| $J_0$ (Liwan District) | 210.465 (36.01,489.293) | $p$ (Liwan District) | 0.011(0.007,0.016) |
| $A_0$ (Liwan District) | 14.164 (12.413,16.164) | $p$ (Nansha District) | 0.027 (0.021,0.034) |
| $p$ (Yuexiu District) | 0.01 (0.006,0.016) | $p$ (Panyu District) | 0.036 (0.027,0.046) |
| $p$ (Haizhu District) | 0.03 (0.022,0.04) | $p$ (Huadu District) | 0.043 (0.03,0.058) |
| $p$ (Huangpu District) | 0.021 (0.016,0.026) | $p$ (Zengcheng District) | 0.045 (0.033,0.06) |
| $p$ (Tianhe District) | 0.026 (0.017,0.041) | $p$ (Conghua District) | 0.004 (0.002,0.007) |
| $p$ (Baiyun District) | 0.017 (0.011,0.024) | | |

**Figure 8:**
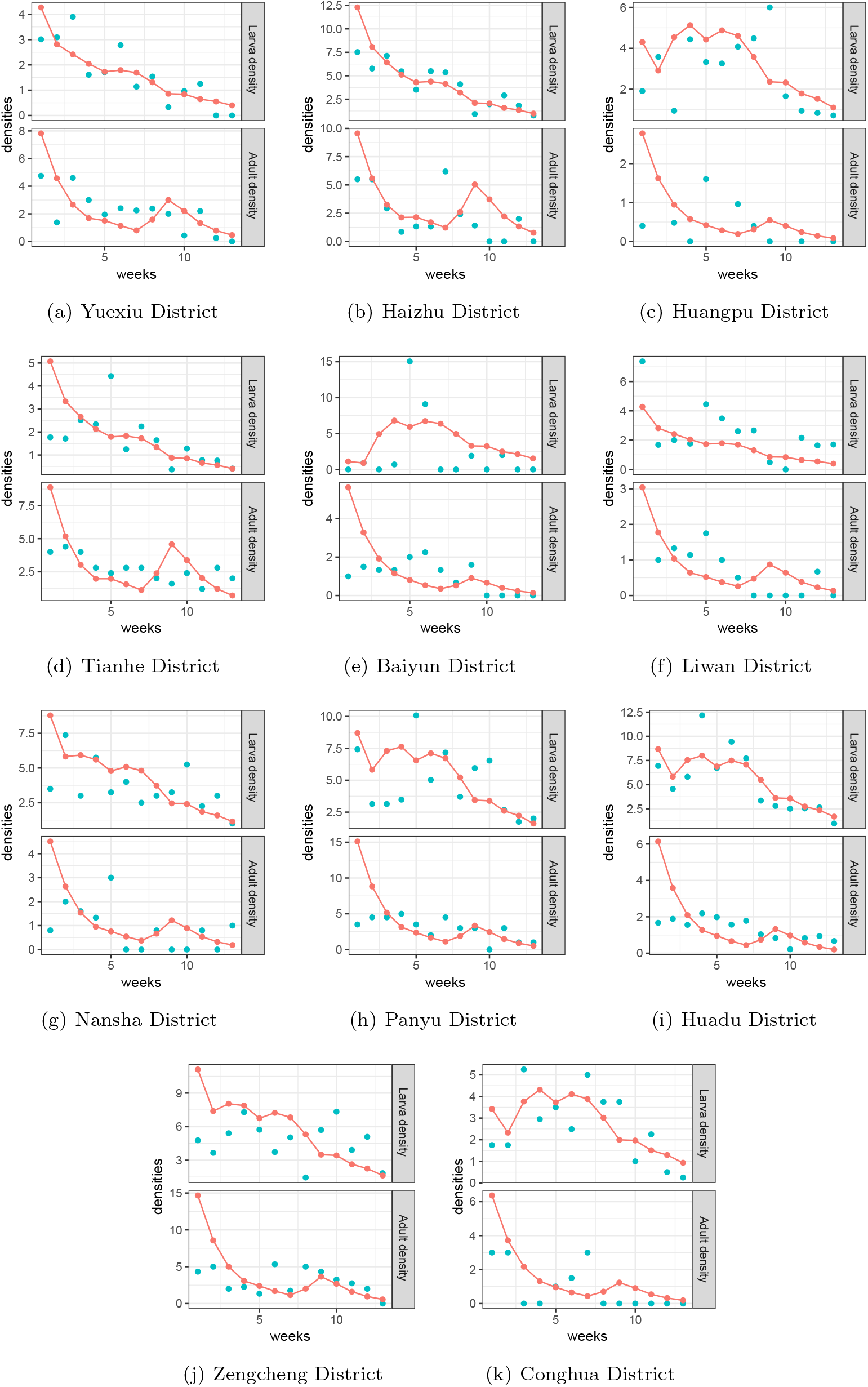
The comparison of the predicted mosquito density (red dots) with the Breteau Index and Bite Index for 11 districts in Guangzhou city in 2017 (blue dots).

**Figure 9:**
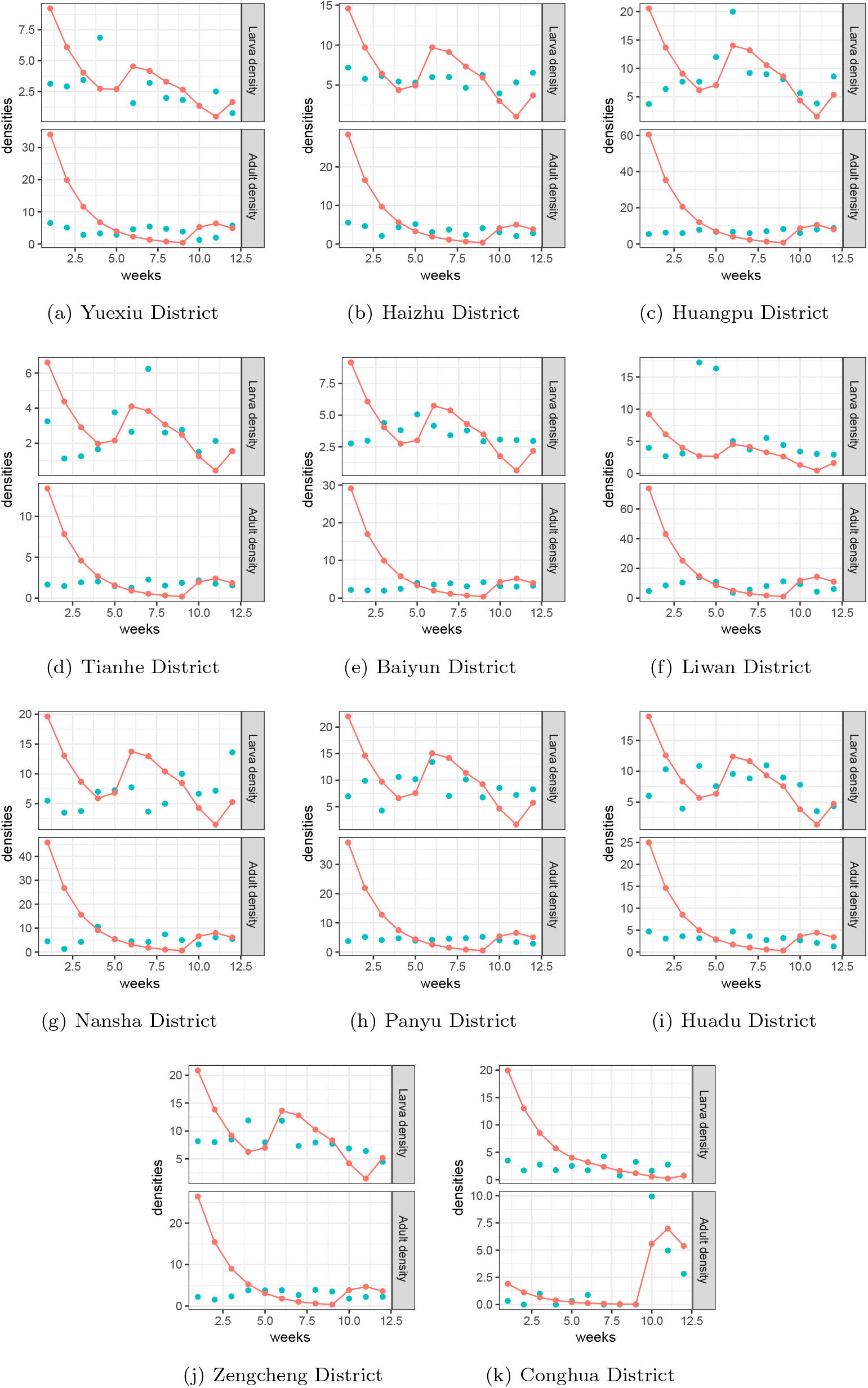
The comparison of the predicted mosquito density (red dots) with the Breteau Index and Bite Index for 11 districts in Guangzhou city in 2018 (blue dots).

## 5 Concluding Remarks

In this paper, we modified the discrete time mosquito model with weather factors developed by Gong et al. (Gong et al., 2011) to describe the mosquito population (specifically Aedes aegypti) in Guangzhou city, China, in 2014. We need to revise the maturation rate as a Guassian function of the temperature, the diapause rate as exponentially decreasing with daylight, and the adult survival rate as temperature independent to identify all model parameters. The resulting new model well agrees with the published Breteau Index and Bite Index for each of the 12 districts in Guangzhou city in 2014.

We also validated our model using the Breteau Index and Bite Index for Guangzhou city in 2017 and 2018, using the model parameters estimated for 2014 except the initial population densities and capture probability ratios, which are fitted independently. The results show well agreement between the model predictions and the data for both 2017 and 2018. Thus, the model and the estimated parameters are suitable for Guangzhou city in other seasons.

The model (1)(a,b) with (3)(a,b,c) may be used to predict the mosquito density in other regions. It is very likely that the model parameters that we estimated in this paper may be applicable to other regions as well, but this needs to be validated using the mosquito data for the regions of interest.

Our results provide a quantification of the weather factors on the mosquito population dynamics. This model can be coupled with disease models to provide a tool for evaluating the risk of mosquito-borne diseases such as dengue infection.

## Acknowledgments

This research was supported by National Natural Science Foundation of China (No.11771075) (ML), Natural Science Foundation of Shanghai (No.21ZR1401000) (ML) and a discovery grant of Natural Sciences and Engineering Research Council Canada (JM).

